# E*x vivo* observation of granulocyte activity during thrombus formation

**DOI:** 10.1101/2020.07.13.199174

**Authors:** Daria S. Morozova, Alexey A. Martyanov, Sergei I. Obydennyi, Julia-Jessica D. Korobkin, Alexey V. Sokolov, Ekaterina V. Shamova, Irina V. Gorudko, Anna Shcherbina, Mikhail A. Panteleev, Anastasia N. Sveshnikova

## Abstract

Infiltration of growing thrombi by leukocytes, being the key part of the thromboinflammation, is well established *in vivo*. The study was aimed at the development of an *ex vivo* simulation of this phenomenon. Thrombus formation in anticoagulated whole blood from healthy volunteers and patients was visualized by fluorescent microscopy in parallel-plate flow chambers with fibrillar collagen type I coverslips.

Moving CD66b-positive cells (granulocytes) were observed in hirudinated or recalcified blood under low wall shear rate conditions (<200 s^−1^). These cells crawled around thrombi in a step-wise manner with an average rate of 70 nm/s. Pre-incubation of blood with leukocyte priming agents lead to a significant increase in average cell velocity. On the contrary, leukocytes from Wiskott-Aldrich syndrome patients demonstrated a 1.5-fold lower average velocity, in line with their impaired actin polymerization.

Thereby, the observed features of granulocytes crawling are consistent with the neutrophil chemotaxis phenomenon. We conclude that the proposed *ex vivo* experimental setting allows us to observe granulocytes activity in near-physiological conditions.

## Introduction

A complex interplay between blood coagulation system, immune system, and endothelium, called thromboinflammation, occurs in diverse pathophysiological situations, such as bacterial infection or cancer (1). Thromboinflammation is thought to be driven mainly by the interactions between neutrophils and platelets (2). Platelets are the circulating in human blood anucleated fragments of megakaryocytes with multiple functions both in hemostasis and immunity (3,4). Platelet activation at the site of injury or inflammation leads to the secretion of platelet α-granules, which contain P-selectin, fibrinogen, vWF, growth factors, and chemoattractants for leukocytes (NAP2, RANTES, CD40L, etc.) (5,6). These proteins play a crucial role in the leukocyte recruitment and adhesion (7–9). The adhesion to platelets causes leukocytes’ integrins activation (10), and their migration through thrombi (11). Therefore, platelet-leukocyte interactions are in the heart of thromboinflammation.

Neutrophils’ migration in thrombi should be dependent on concentrations of their chemoattrants. Contact with a chemotractant (“priming”) of neutrophils leads to their β2-integrins (CD11a/CD18, LFA-1 (αLβ2) and CD11b/CD18, Mac-1 (αMβ2)) activation, which results in their firm adhesion to the surface. Therefore, we expect that neutrophil movement in a thrombus should be influenced by priming. An inhibition of actin polymerisation impairs neutrophil chemotaxis (12). In patients with cytoskeletal abnormalities, for instance, with Wiskott-Aldrich syndrome (genetic hemorrhagic and immunological syndrome; WAS) (13,14), we expect impaired neutrophil involvement in the thrombus formation process.

The study was aimed at the development and validation of an *ex vivo* technique, allowing observation and simulation of the thrombus-leukocyte interactions (thrombus growth and leukocyte activity). To validate the method we loaded the hirudinated (anticoagulated) whole blood of healthy donors or WAS patients in parallel-plate flow chambers under the low flow shear rate (100 s^−1^). The samples from healthy donors were studied under control conditions as well as with leukocyte-priming reagents. We have identified conditions for granulocytes’ observation and derived a plethora of parameters for granulocytes’ characterization. These parameters were used for the analysis of the whole blood of WAS patients.

## 2. Materials and Methods

### 2.1. Materials

The sources of the materials were as follows: Annexin V-Alexa Fluor 647 (BioLegend, San Diego, CA), DiOC-6, Fucoidan from Fucus vesiculosis, HEPES, bovine serum albumin, lipopolysaccharides from E. coli O111:B4, human fibrinogen (Sigma-Aldrich, St Louis, MO); CD11b-FITC, CD66b-PE (Sony Biotechnology, San Jose, CA), fibrillar collagen type I (Chrono-Log Corporation; Havertown; USA);. Isolated from the milk of transgenic goats, lactoferrin (rLf) was kindly provided by Semak I.V. (Department of Biochemistry, Biological Faculty, Belarusian State University, Minsk, Belarus) (15).; human von Willebrand Factor (vWF) was a kind gift of Prof. Pierre Mangin (NSERM, Etablissement Français du Sang-Grand Est, UMR_S1255, Fédération de Médecine Translationnelle de Strasbourg, Université de Strasbourg, France). The HL-60 cell line (promyelocytic leukemia) was used as a source of myeloperoxidase (MPO). MPO was isolated, as described in (16).

### 2.2. Blood collection and handling

Blood collection was performed under the protocol approved by the free CTP PCP RAS Ethical Committee (protocol #1 from 12.01.2018), and written informed consents were obtained from all donors and patients. Blood was collected from healthy adult volunteers (n=30, men and women 18-35 years old) into Vacuette© sodium citrate (3.8 % v/v) or lithium heparin (18 I.U./ml blood) or Sarstedt-Monovette© hirudin (525 ATU/ml blood) vacuum tubes. Experiments were performed within 3 hours after blood collection. For the assays involving Wiskott-Aldrich syndrome patients, blood was collected from healthy pediatric donors (n = 10) or from patients with Wiskott-Aldrich syndrome (n = 7) into Sarstedt-Monovette hirudin (525 ATU/ml blood) tubes. None of the patients received specific therapy at the time of the experiments.

Blood samples were purified from plasma proteins by 3 sequential centrifugations of citrated whole blood for 10 minutes by 1000g with supplementation of the plasma by Tyrodes’s calcium-free buffer. Final supplementation was performed by Tyrodes’s calcium buffer.

### 2.3. Fluorescent microscopy

Parallel-plate flow chambers were described previously (17). Channel parameters were: 0.2 x 18 x 0.206 mm. Glass coverslips were coated with fibrillar collagen type I (0.2 mg/ml) for 1h 30min at 37°C, washed with distilled water and then inserted into the flow chambers.

Whole blood was pre-loaded with DiOC6 (1 μM) and AnnexinV-Alexa647 (10 μg/mL). Blood was perfused through the parallel-plate chambers over collagen-coated (0.2 mg/ml) surface with wall shear rates 100 s^−1^ (18). Thrombus growth and leukocyte crawling were observed in DIC/epifluorescence modes with an inverted Nikon Eclipse Ti-E microscope (100x/1.49 NA TIRF oil objective). Alternatively, confocal mode with Z1 microscope (Carl Zeiss, Jena, Germany; 100x objective; Axio Observer) was used.

### 2.4. Data analysis

Nikon NIS-Elements software was used for microscope image acquisition; ImageJ (http://imagej.net/ImageJ) was used for image processing. ImageJ manual tracking plugin was used for manual granulocyte tracking, and the Coloc2 plugin for fluorescence colocalization analysis plugin was utilized.

For automated cell tracking, particle tracking algorithm described in (19) was utilized. The algorithm was based on Python trackpy v.0.4.2 library. First, particle tracking was performed, then the tracks belonging to leukocytes were selected manually. The platelet thrombus area was calculated as the percentage of the screen covered by platelet thrombi upon the subtraction of the area of crawling cells. Tracking Code listing and program operation examples can be found in Supplementary Materials.

### 2.5. Statistics

All experiments were performed at least in triplicate with platelets from different donors. Statistical analysis was performed using Python 3.6; all statistical details are provided in the figure captions.

## 3. Results

### 3.1. Granulocytes crawl among the growing thrombi under low shear rate in the presence of calcium in the whole blood

Collagen coated parallel plate flow chambers are a widely applied modern tool for the hemostasis assessment (20). They are mostly used to mimic micro-vessels, such as arterioles (high blood flow velocity) and post-capillary venules (low blood flow velocity) (21). As has been demonstrated *in vivo*, immune cells participate predominantly in the venule hemostasis (22), thus, here we aimed to reconstruct the venule conditions *ex vivo*. Under the low shear rate conditions (100 s^−1^) in sodium citrate-anticoagulated (plasma coagulation was inhibited by chelating calcium and magnesium ions) whole blood, normal platelet thrombus growth, without inclusions of other cell types, was observed (Fig. 1A-C). On the other hand, in hirudin-anticoagulated whole blood (anticoagulation effect was achieved by direct thrombin inhibition) inclusions of cells with diffuse DiOC-6 distribution was observed (Fig. 1D-F). These cells rolled over, adhered, and began crawling among the growing platelet thrombi (Supplementary video 1). These crawling cells were CD66b (Fig. 1G) and CD11b (Fig. 1H). Therefore, we claim that these cells were granulocytes. The same phenomenon was observed in the heparin-anticoagulated or citrate-anticoagulated blood upon recalcification (Fig. S1A-F), which emphasized the role of calcium ions in the granulocyte involvement in the hemostatic processes under low shear rate. Hirudin- and heparin-anticoagulated blood was used in all further experiments because citrated blood recalcification causes local fibrin formation and platelet activation (22).

**Figure 1.**
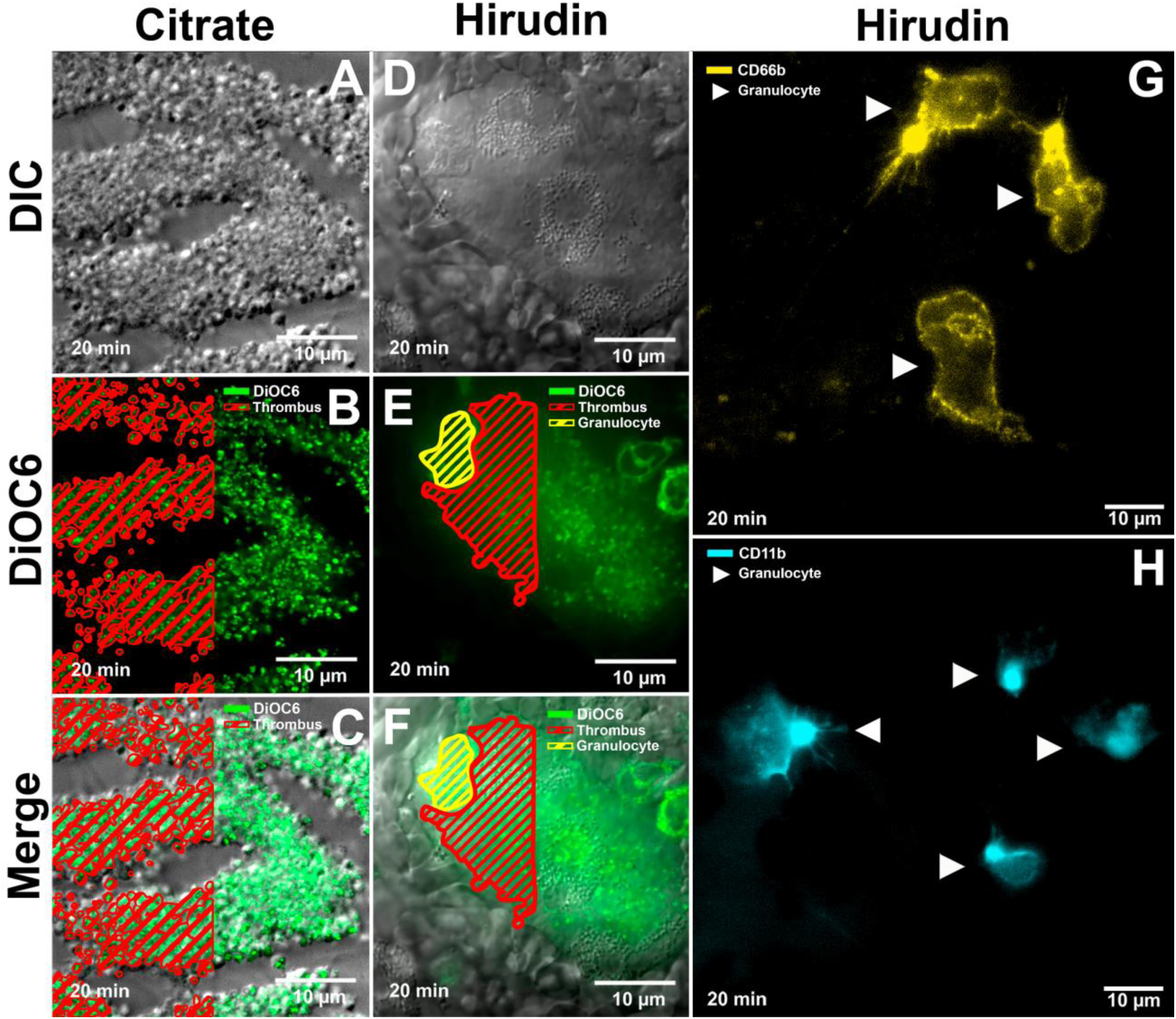
Thrombus growth and immune cell behavior in citrated or hirudinated blood. (A-C) Citrate- or (D-F) hirudin-anticoagulated blood was perfused over collagen covered coverslips under the shear rate of 100 s-1. Growing thrombi were visualized in DIC (A,D) or in confocal mode upon DiOC6 staining (green, B,E). Merged images are given in C,F. Platelet thrombi are indicated by red (half of the thrombi is highlighted). Crawling cells with diffuse DiOC6 staining are indicated by yellow (half of each cell is highlighted). (G,H) Crawling cells appeared to be CD66b (G) and CD11b (H) positive. Representative data out of n = 5 experiments.

### 3.2. Granulocytes behavior change in the process of crawling among the growing thrombi

Despite the significant diversity in the crawling granulocytes’ behavior, two major types of the crawling cells could be identified – granulocytes with uniform DiOC6 staining (“type A”, Fig. 2A) and granulocytes with cluster-like DiOC6 staining (“type B”, Fig. 2B). Observation of the fate of each cell, specifically (Fig. 2C, Supplementary video 2) allowed us to conclude that in the process of crawling, granulocytes underwent the transition from type A (Fig. 2A) to type B (Fig. 2B). During this process, cells slowed down (velocity < 0.045 µm/min, Fig. 2D) and spread (Fig. 2B,C, Supplementary video 2). This behavior allows us to suppose that the “type B” cells are more activated than “type A”.

**Figure 2.**
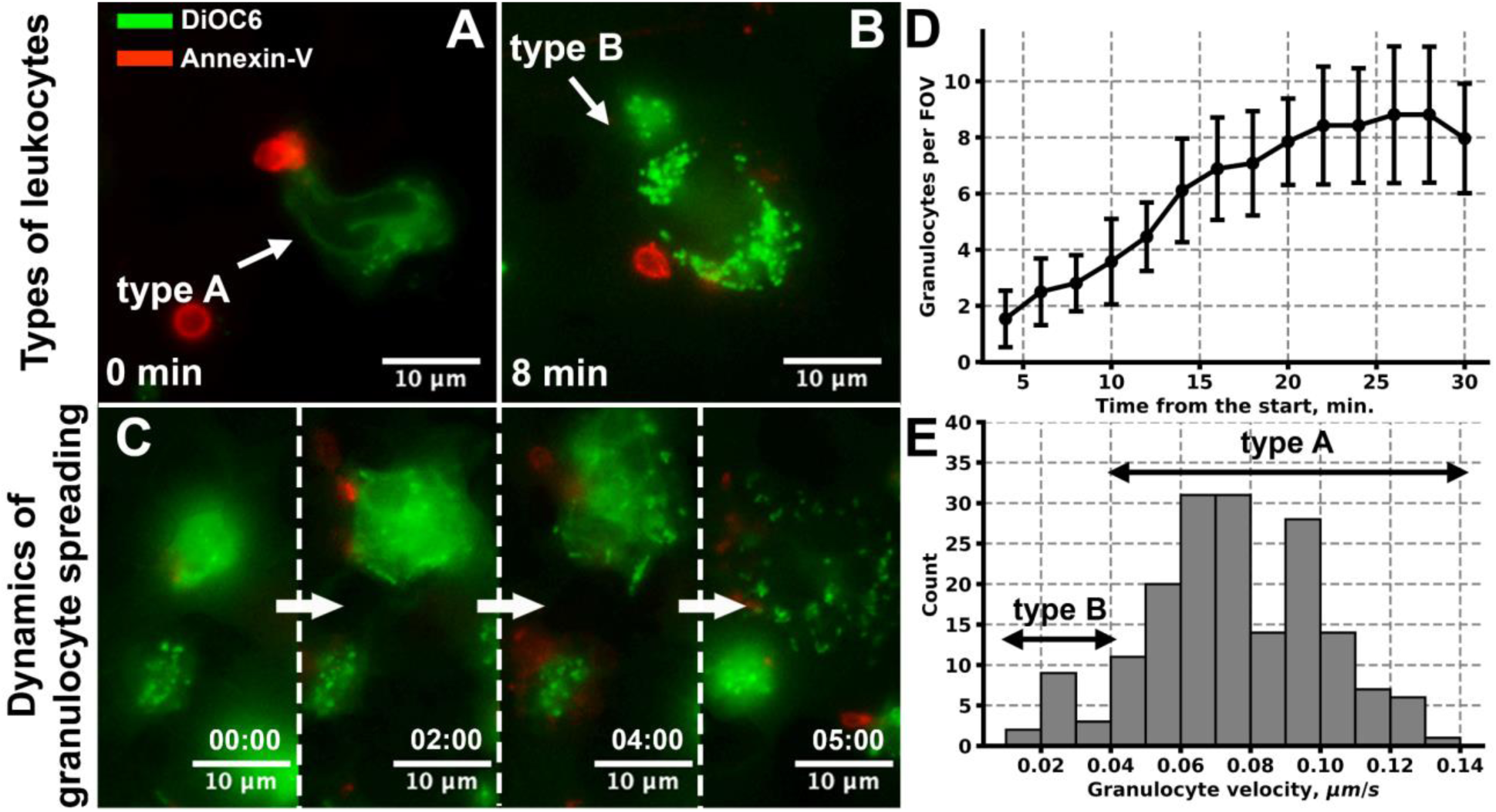
Crawling granulocytes behavior. (A-C) Crawling granulocytes formed two types: cells with diffuse DiOC6 staining (type A, A) and cells with cluster-like DiOC6 staining (type B, B). With the flow of time, the transition from type A to type B occurred (C). (D) The number of granulocytes per FOV increased gradually up to the 20^th^ minute of the experiment. (E) Based on the granulocyte velocity distribution, it can be claimed that type B granulocytes were distinctive not only by their appearance but by their velocity as well. Representative data out of N = 10 donors.

Under the described conditions, the number of granulocytes gradually increased from 1.6 cells per field of view (FOV) on the 5^th^ minute to 8 cells per FOV on the 20^th^ minute and did not change significantly up to 30^th^ minute (Fig. 2E). A comparison between heparin- and hirudin-anticoagulated blood revealed that the total number of granulocytes per FOV was higher in the heparinized blood (Fig. S1G). Numbers of type B granulocytes did not differ significantly in heparin- or hirudin anticoagulated blood (Fig. S1H). This allowed us to claim, in agreement with previously published data (23), that heparin weakly pre-activates granulocytes in our experimental setting. Thus, hirudin-treated blood samples were used in all further experiments.

### 3.3. Mediators of inflammation and platelet activators alter granulocyte behavior among the growing thrombi

Motile type A granulocytes crawled on collagen among the growing thrombi independently of the blood flow (Fig. 3A,B), following a curved trajectory (Fig. 3C). Velocity changed from 0.014 to 0.21 µm/s (Fig. 3C), with the mean velocity of 0.07±0.01 µm/s (Fig. 3G). Upon instant velocity averaging over time, granulocyte movement appeared to be more uniform (Fig. S2). Thus, it can be claimed that a direct granulocyte movement is observed.

**Figure 3.**
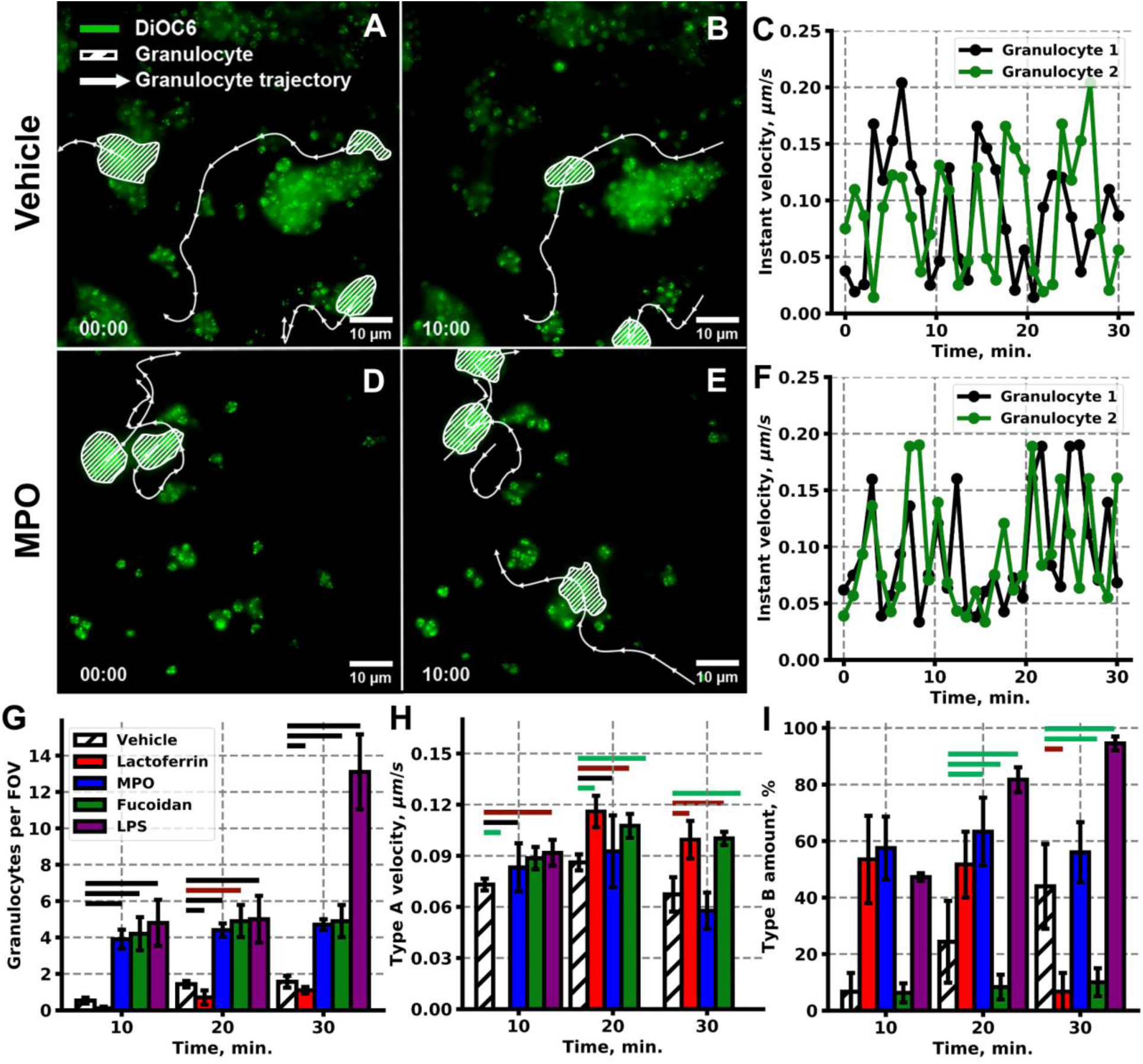
Granulocyte and platelet activators impact on granulocyte behavior among the growing thrombi. (A-C) Granulocytes from vehicle-treated blood crawl among platelet thrombi in a direct manner: A – initial moment, B – 10 minutes after granulocyte adhesion. (C) Instant granulocyte velocity appeared to be intermittent, although averaged velocity was uniform. (D-F) Granulocytes from MPO treated blood crawl among platelet thrombi in a direct manner: D – initial moment, E – 10 minutes after granulocyte adhesion. (F) Instant granulocyte velocity appeared to be intermittent, although averaged velocity was uniform. (G-I) Granulocyte and platelet activators alter the number of granulocytes per FOV (G), type A granulocyte velocity (H), and type B granulocyte amount (I). Statistical significance was calculated with the Mann-Whitney test; green lines correspond to p < 0.05, red lines correspond p < 0.01, black lines correspond p < 0.001. Statistics were calculated over 20 FOVs from n = 10 donors.

Our original hypothesis was that granulocytes adhered to the growing thrombi in an activation-dependent manner. Thus, we performed experiments in the presence of leukocyte-activating agents: myeloperoxidase (MPO; (24)), Lactoferrin (25) and lipopolysaccharides (LPS; (26)) to mimic the pro-inflammatory granulocyte stimulation or fucoidan (27) for simultaneous weak activation of both granulocytes and platelets (Supplementary video 3). MPO (Fig. 3D-F) and Fucoidan (Fig. S3G-I) did not significantly affect granulocyte trajectories, while LPS and Lactoferrin decreased the length of granulocyte trajectory (Fig. 3A-F). All activators, except for Lactoferrin, significantly increased the number of granulocytes per FOV at all time points (Fig. 3G). Pre-incubation with MPO and Fucoidan resulted in the statistically significant increase of the mean granulocyte velocity at all time points (Fig. 3H, blue and green bars, correspondingly). Lactoferrin reduced granulocyte number to 0 at 10^th^ minute, but increased granulocyte velocity at 20^th^ and 30^th^ minutes of the experiment (Fig. 3H, red bars). On the other hand, LPS increased granulocyte velocity only at 10^th^ minute but significantly reduced at 20^th^ and 30^th^ minute (Fig. 3H, purple bars), which was associated with the statistically significant increase of the numbers of slow spread type B granulocytes upon LPS treatment (Fig. 3I). MPO and Lactoferrin also increased numbers of type B granulocytes, although statistical significance was not reached (Fig. 3I). None of the agonists, except for LPS, altered thrombus growth at the described conditions (Fig. S4A).

### 3.4. Crawling granulocytes bear Annexin-V positive platelets

Removal of the phosphatidyl-serine (PS) exposing (Annexin-V – positive) cells is one of the key physiological functions of granulocytes in the blood flow (28). Upon strong stimulation, platelets expose PS on their outer membrane and become procoagulant instead of being pro-aggregate (29). This results in platelet extrusion from the core of the growing thrombi (17), where they can be picked up by the crawling granulocyte.

Granulocytes associated with Annexin-V positive platelets were observed in our *ex vivo* conditions (Fig. 4A-F, S5A-C). However, colocalization analysis (30) revealed that DiOC-6 (Fig. 4G) and CD66b (Fig. S5D) staining did not significantly correlate (Pearson’s correlation coefficient, PCC, 0.46±0.13 and 0.52±0.15, correspondingly) with Annexin-V fluorescence (Fig. 4I). On the other hand, the correlation between CD11b and Annexin-V fluorescence (Fig. 4H) was significantly higher, PCC = 0.68±0.12 (Fig. 4I). Based on this finding, it can be claimed that Annexin-V labeled platelets attach to crawling granulocytes via CD11b. Further analysis of the behavior of the Annexin-V bearing DiOC6 stained granulocyte revealed that, despite low dye correlation (Fig. 4G,I), trajectories of the crawling granulocytes and Annexin-V labeled granulocyte (Fig. 4J-L, S5E) corresponded in all cases.

**Figure 4.**
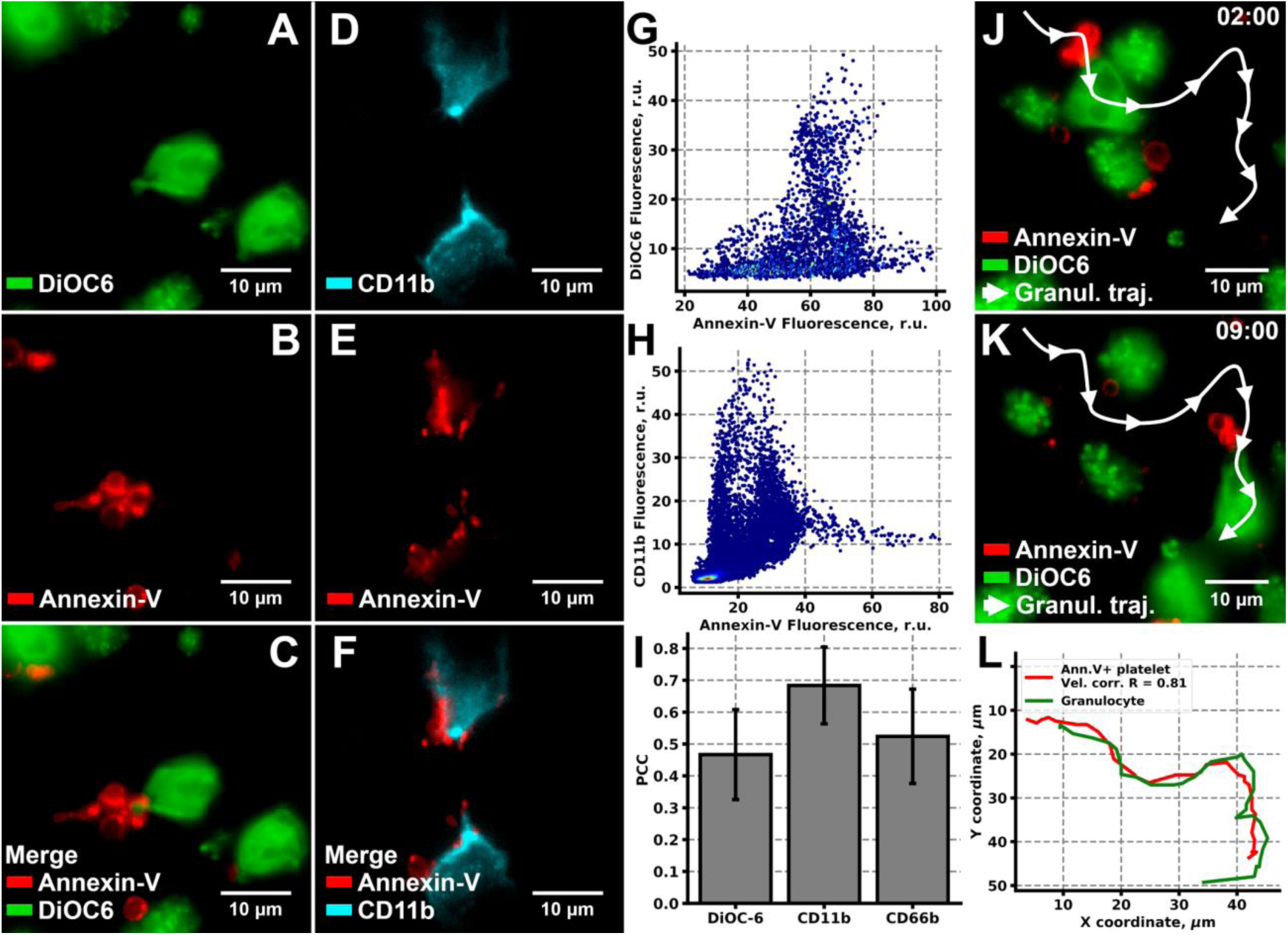
Annexin-V positive platelets attach to the crawling granulocytes. (A-C) DiOC6 (A) and Annexin (B) staining colocalized (C) in a manner suggesting that granulocytes bear Annexin-V positive platelets. (D-F) Similar conclusions can be made based on CD11b (D) and Annexin-V (E) colocalization (F) analysis. (G,H) Scatterplots of the Annexin-V and DiOC6 (G) or CD11b (H) fluorescence intensity correlation from (C) and (F), correspondingly. (I) Pearson’s R correlation coefficient (PCC) for Annexin-V and DiOC6, CD11b and CD66b dyes (n = 50 for each pair). (J-K) Microscopy images of the crawling granulocyte, bearing Annexin-V positive platelets after 2 minutes (J) and after 9 minutes (K) from the start of the observation. (L) trajectories of the granulocyte (green curve) and the Annexin-V positive platelet (red curve) from (J-K). Typical results out on n = 50.

### 3.5. Plasma proteins and calcium ions are required for granulocyte crawling among the thrombi

Previously we have demonstrated that granulocytes’ incorporation into the growing thrombi requires free calcium ions (Fig. 1). In order to investigate the mechanisms of the granulocyte involvement in the platelet thrombus formation process, we have removed plasma proteins from the whole blood by means of sequential centrifugation (see Methods). In such a “clear” system, no granulocyte crawling on the collagen surface was observed (Fig. 5A), while short-term granulocyte attachment to the platelet covered surface and granulocyte rolling was observed 3.6 µm above the collagen layer (Fig. 5B, Supplementary video 4). The introduction of 10% of physiological concentrations of fibrinogen and von Willebrand Factor (vWF) resulted in the recovery of granulocyte crawling (Fig. 5CD) on the collagen covered glass, yet granulocytes descended from the thrombi less readily than in the whole blood (Fig. 5EF). Based on these findings, we propose the following scheme of the events: granulocytes from the blood flow attach to the growing thrombi in the presence of calcium and descend onto the collagen level via fibrinogen and vWF, where granulocytes crawl (Type A), bearing Annexin-V positive platelets and eventually slow down and spread (Type B, Fig. 5G).

**Figure 5.**
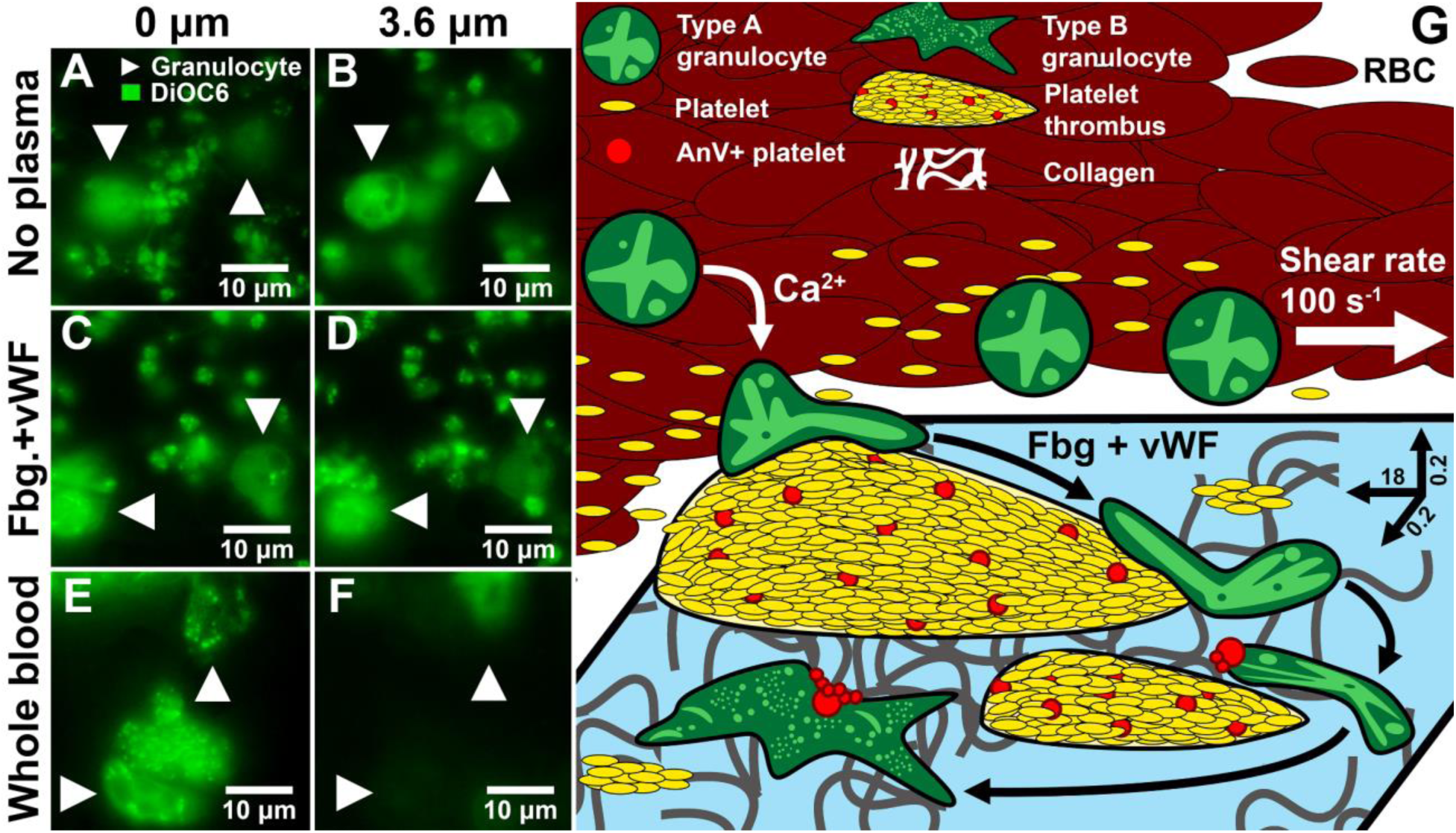
The role of blood plasma proteins in granulocytes’ incorporation into the growing thrombi. (A-F) Confocal microscopy images of the growing thrombi and granulocytes. (A,B) Thrombi in the “clean” system on the collagen level (A) and 3.6 µm above (B). (C,D) Thrombi in the “clean” system with the addition of fibrinogen and vWF on the collagen level (C) and 3.6 µm above (D). (E,F) Thrombi in the whole blood on the collagen level (E) and 3.6 µm above (F). Representative images of n = 10 experiments. (G) Scheme of the granulocyte behavior on the collagen covered glass coverslip (in 0.2×0.2×18 mm flow chamber): granulocytes (green cells) attach to the growing thrombi (yellow) in the presence of calcium ions. In the presence of fibrinogen and vWF granulocytes, descend to the collagen level and collect Annexin-V positive platelets from the thrombi in the process of crawling (Type A). Eventually, granulocytes slow down and spread on the collagen (Type B).

### 3.6. Crawling of WAS patients’ granulocytes is altered in comparison to healthy donors

Based on our previous findings, we assumed that granulocytes’ behavior in the growing thrombi would be altered in a case of a disease with abnormal cytoskeletal functions. Wiskott Aldrich syndrome is such a genetic disease, in which blood cells cytoskeletal defects have been described as a result of the *WAS* gene mutations (rev. in (31)). WAS is mainly characterized by immunodeficiency, microthrombocytopenia, and autoimmune/oncological predisposition (rev. in (14)).

Typical FOVs of healthy donors and WAS patients are shown in Fig. 6 (A-C and D-F, correspondingly). Qualitative analysis revealed that thrombus areas of WAS samples were significantly reduced (Fig. S4B), while the number of granulocytes per FOV was increased (Fig. S4C) in comparison with control samples. This resulted in a significantly increased ratio of granulocyte number to thrombus area in WAS samples (Fig. 6G). On the other hand, type A granulocytes of WAS patients were significantly slower than type A granulocytes of healthy donors (Fig. 6G). Finally, type B granulocytes amount was significantly increased in WAS samples in comparison with control samples (Fig. 6H). Based on this, it can be claimed WAS granulocytes are more prone to the collagen adhesion and spreading, possibly due to their increased reactivity.

**Figure 6.**
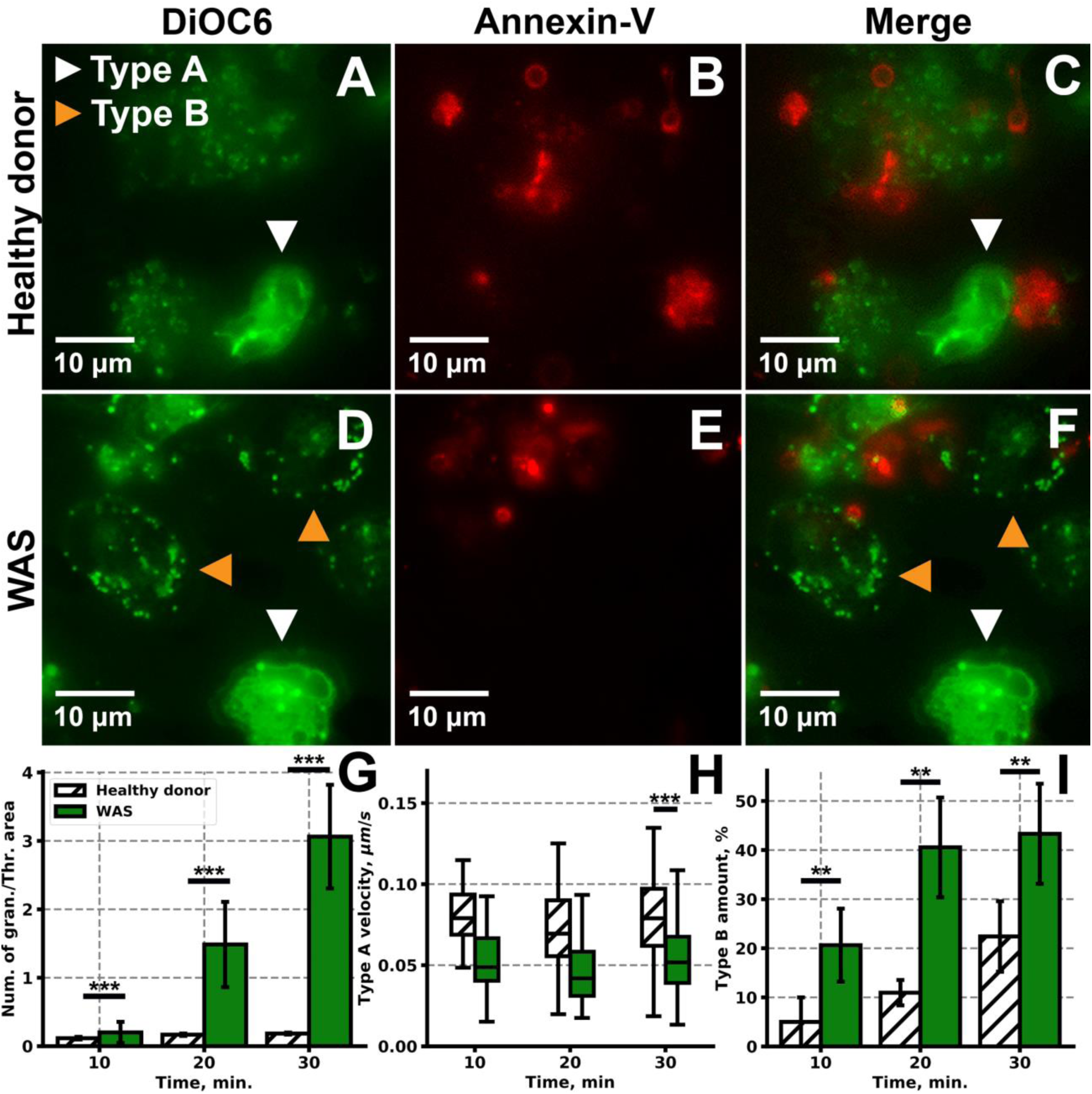
Comparison of healthy donors’ and WAS patients’ granulocytes behavior. (A-C) Typical for healthy donors FOV with growing thrombi and crawling granulocytes. (D-F) Typical for healthy donors FOV with growing thrombi and crawling granulocytes. (G-I) Quantitative comparison of the granulocyte behavior of healthy donors (white) and WAS patients (green): the ratio of granulocyte number to thrombus area (G), average velocities of crawling of type A granulocytes (H), amounts of slow spread type B granulocytes (I) for n = 10 healthy donors and n = 7 WAS patients. Statistical significance was calculated with Mann-Witney test, ** corresponds to p < 0.01, *** corresponds to p < 0.001.

## Discussion

*Ex vivo* approaches to the characterization of the interplay between the hemostasis and immune systems have been limited, though attempts to characterize granulocyte behavior have been previously made (7,8,22). Here we utilized parallel-plate fibrillar collagen-coated flow chambers and low shear rate (100 s^−1^) to mimic venule conditions. We observed granulocyte rolling, arrest, crawling, and spreading (Fig. 1,2) in an activation-dependent manner (Fig. 3). Most granulocytes were bearing Annexin-V positive platelets, and it can be claimed that this interaction was mediated by granulocyte CD11b (Fig. 4). Granulocyte involvement in the thrombus growth appeared to be mediated by plasma proteins – fibrinogen and vWF (Fig. 5). Therefore, several features, characteristic for the neutrophil chemotaxis phenomenon (32), were observed here. The overall pattern of leukocyte participation in thrombi growth *in vivo* correlated with the results obtained *ex vivo* in our study. Lower crawling velocities and the higher relative amount of granulocytes were observed in WAS patients’ samples (Fig. 6). The exact meaning of these results for WAS pathology is unknown, yet they demonstrate that granulocyte-thrombi interaction depends on the intact cytoskeleton function. Also, these and other findings of the study suggest that the proposed experimental settings can be utilized for the *ex vivo* observation and investigation of thromboinflammation.

The first quantitative characteristic of granulocytes proposed here is their movement velocity, which appears to be 0.07±0.02 µm/s on the average, with instant velocities reaching 0.2 µm/s (Fig. 3). These values increased upon priming of neutrophils with various agents (Fig. 3) and decreased in WAS samples (Fig. 6). Together these features support the conclusion that the observed movement of granulocytes around thrombi is the chemotaxis phenomenon (32,33). Previously, *C. Jones et al*. showed that human neutrophils migrated towards LTB4 with an average velocity of 0.39±0.09 µm/s (32). While *Jones et al*. used a constant gradient of chemoattractant, in our experimental setting there were several target areas of attraction for neutrophils, which might explain non-monotonous movements of granulocytes in the current study (Fig. 3). In another study, *M. Weckmann et al*. observed the velocity of neutrophil migration on fibronectin in the presence of IL-8, fMLP, and LTB4 (33), where leukocyte velocity appeared to be 0.11±0.12 µm/s, the same as in our study in the presence of the priming reagents (Fig. 3).

The second quantitative charachteristic observed here is the persentage of activated granulocytes. It is well known that neutrophils’ interactions with platelets during thrombotic events can lead either to phagocytosis or NETosis, depending on the activation triggers (34). Unconventional or longstanding sterile stimuli result in suicidal vs. vital NET generation both of which become the final irreversible stage of leukocyte activation, causing their immobile state (35,36). In the current study we created sterile pro-inflammatory settings with gradual thrombus growth, that resulted in platelet activation and dying. Thus we conclude that the slowly moving or immobile granulocytes with poorly DiOC6-stained cytoplasm are the highly activated cells.

The third semi-quantitative characteristic is the total number of granulocytes per field of view (Fig. 3). Obviously, this number depends on the concentration of platelets, because it is well-known that platelets activate and attract leukocytes by platelet-leukocyte interactions during thromboinflammation (1,9,37). However, the influence of leukocytes on platelet activation and thrombus growth is under-investigated. Here, in line with previous studies (25,38–43), granulocytes’ activation resulted in granulocyte shape and velocity changes (Fig. 3G-I), as well as in the changes in thrombus area (Fig S4A). This additionally allows us to claim that granulocyte and platelet activation could be monitored using the proposed experimental settings.

Altogether, the proposed experimental settings can be used for the assessment of platelet-neutrophil interplay in different conditions, including hematological and immunological disorders, such as WAS. Here, using the proposed method, we demonstrate that the average velocity of granulocytes’ crawling is significantly lower in WAS, while the number of attracted leukocytes is significantly higher than those for healthy donors (Fig. 6). Migration of granulocytes was shown to be impaired in WAS KO mice (44). In WAS patients granulocyte migration defect appears to be more subtle and dependent on the integrin signalling (45,46). The WASP-deficient neutrophils also showed reduced integrin-dependent degranulation and respiratory burst (47). On the other hand, enhanced granulocyte recruitment to the growing thrombi, observed in our study, can be the result of the higher proportion of procoagulant platelets in WAS patients (48,49). Therefore, the findings of our study are consistent with previous reports on platelets and granulocytes in WAS.

Several other new findings of platelet-granulocyte interplay were observed in our study. First, we observed procoagulant platelets attached to the moving granulocytes (Fig. 4). Annexin-V positive, procoagulant platelets on the margins of thrombi cause thrombin generation, affecting blood plasma coagulation (17). Procoagulant platelets in our conditions attracted granulocytes in the manner common for any tissue debris (Fig. 4), as demonstrated in several studies (50). Fluorescent dye correlation analysis results (Fig. 4D-F, S5) allow us to claim that CD11b mediates this interaction. We suggest that one of the functions of granulocytes in thrombus formation is the scavenging of procoagulant platelets (Fig. 4G-I, S5E) and thus, limiting thrombus growth in venules. Second, we propose that the neutrophil involvement in thrombus formation is dependent on plasma proteins (fibrinogen and vWF) (Fig. 5A-F), in line with findings of *Constantinescu-Bercu A. et al*.. (51) who demonstrated vWF- and platelet integrin αIIbβ_3_ –dependendent activation of neutrophils and with findings of *Ghasemzadeh M. et al*. (22), who confirm the essential role of fibrin in intravascular leukocyte trafficking. Additionally, the vWF role in the recruitment of leukocytes during thromboinflammation has been previously established (52). However, the exact mechanism of granulocyte crawling on collagen has to be the object of further studies.

In this study, we report the phenomenon of the granulocyte crawling among the growing thrombi *ex vivo*. We claim that granulocyte characterization can be used for a more in-depth analysis of the mechanisms of immunological and hematologic diseases. Based on our own experimental assays, we propose a scheme of granulocyte participation in thrombus formation: (1) granulocytes attach to growing thrombi in a calcium-dependent manner; (2) granulocyte descent to collagen level is mediated by fibrinogen and vWF; (3) descended granulocytes collect annexin-V positive platelets from the growing thrombi; (4) granulocytes eventually slow down and spread.

## Authors’ contributions

D.S.M. performed fluorescent microscopy experiments, analyzed the data and wrote the paper; A.A.M. performed fluorescent microscopy experiments, analyzed the data and edited the paper; S.I.O. performed confocal microscopy experiments; J-J.K. developed the software for the automated data analysis; A.V.S. isolated MPO and edited the paper; E.V.S. and I.V.G. analyzed the data and edited the paper; A.Y.S. managed the patients with WAS and edited the paper; M.A.P. planned the research and edited the paper; A.N.S. supervised the project, planned the research, analyzed the data, performed experiments and edited the paper. The authors declare that they have no conflict of interest.

## Acknowledgments

We thank Miss. A.E. Boldova for her assistance during the microscopy data collection.

## Funding

The project was supported by a grant from the endowment foundation “Doctors, Innovations, Science for Children”, and by the Russian Foundation for Basic Research Grant 17-00-00138.

## Abbreviations

MPO: myeloperoxidase
rLf: recombinant lactoferrin
LPS: lipopolysaccharides from E. coli O111:B4
vWF: von Willebrand factor
PS: phosphatidylserine
GP: glycoprotein
CD: cluster of differentiation
FOV: field of view
WAS: Wiskott-Aldrich syndrome.

## Supporting Information

**Figure S1.**
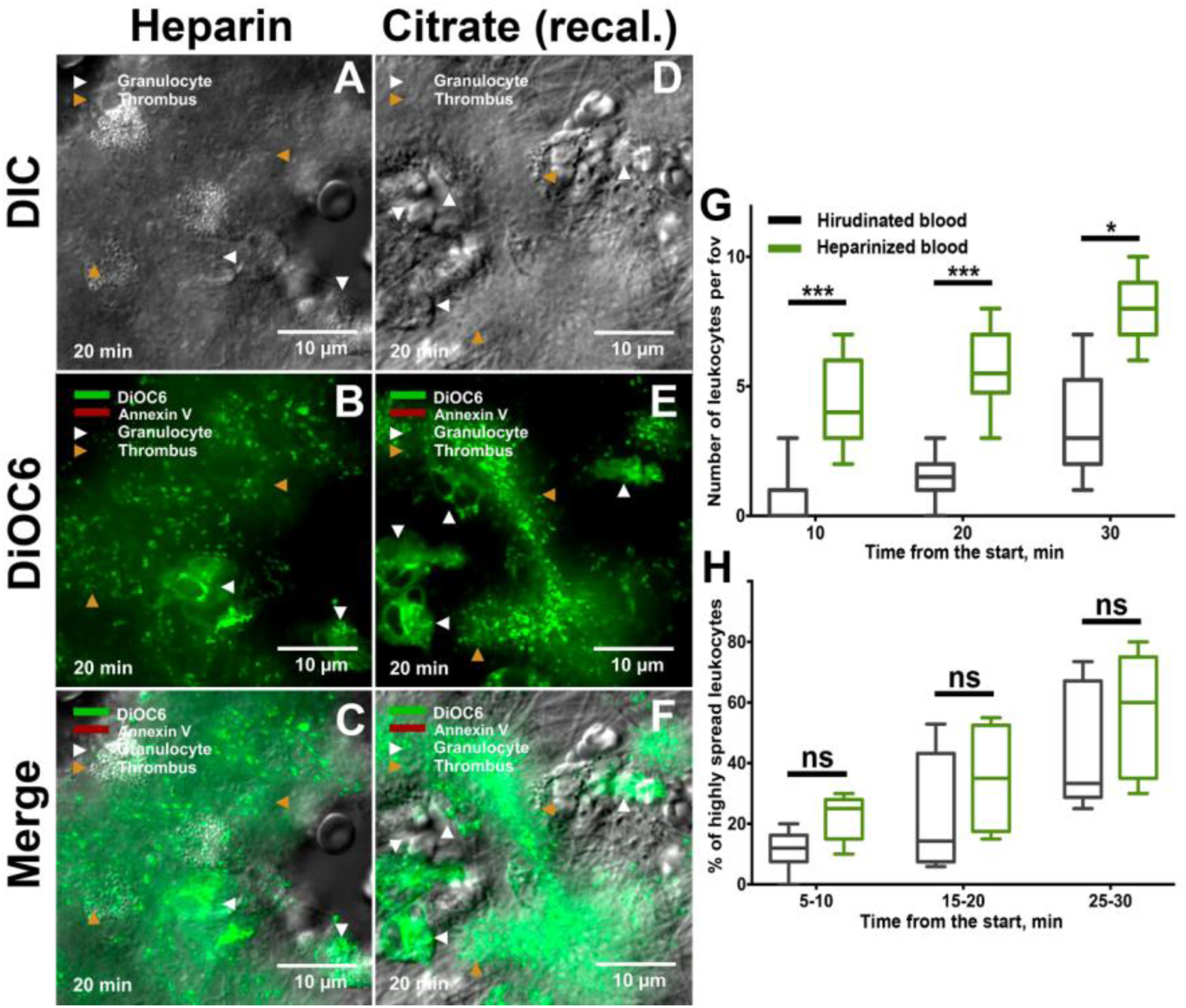
Conditions for leukocyte observation in parallel-plate flow chambers. Anticoagulated whole blood was perfused through parallel-plate flow chambers above fibrillar collagen type I surface with the permanent wall-shear rate of 100 s^−1^. Confocal mode was used to visualize thrombi and leukocytes in DIC; DiOC6 stain was applied in order to visualize cell membranes. Images presented were captured on the 20^th^ minute of perfusion. Orange triangles present thrombi, white – leukocytes; from left to right: DIC mode; DiOC6 staining (green); merged. A – citrated blood upon recalcification; D – heparinazed blood. (G) –comparison of granulocyte number in heparinazed and hirudinated blood; (H) – percentage of highly spread granulocytes in whole blood with different anticoagulants at different time intervals. Statistical significance was calculated with Mann-Wittney test, asterisk (*) indicates p = 0.01.

**Figure S2.**
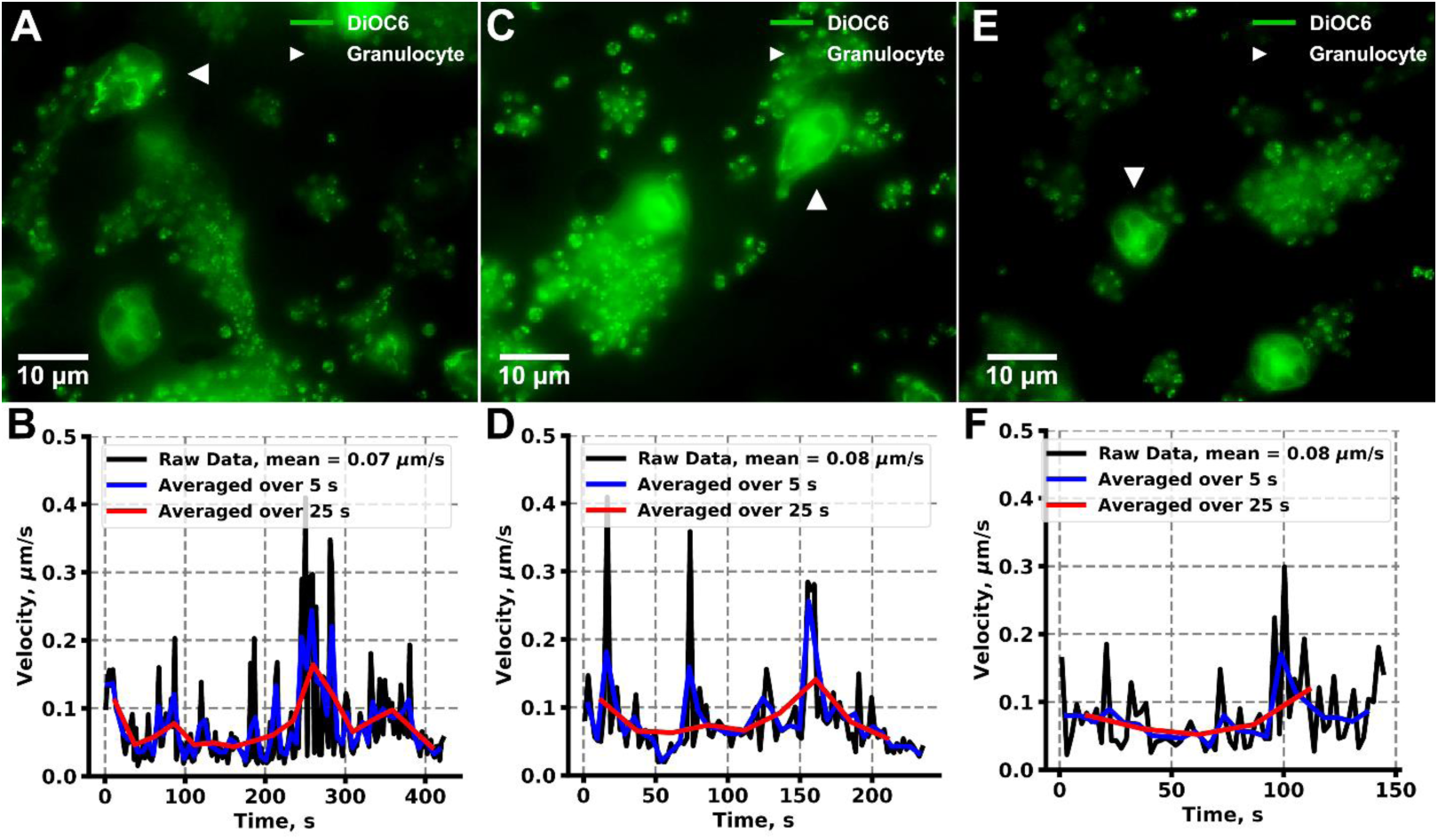
Instant leukocyte velocities and averaged velocities. Epifluorescence (DiOC6 stain – green) mode was used. Images and corresponding velocity of healthy different granulocytes of a healthy donor.

**Figure S3.**
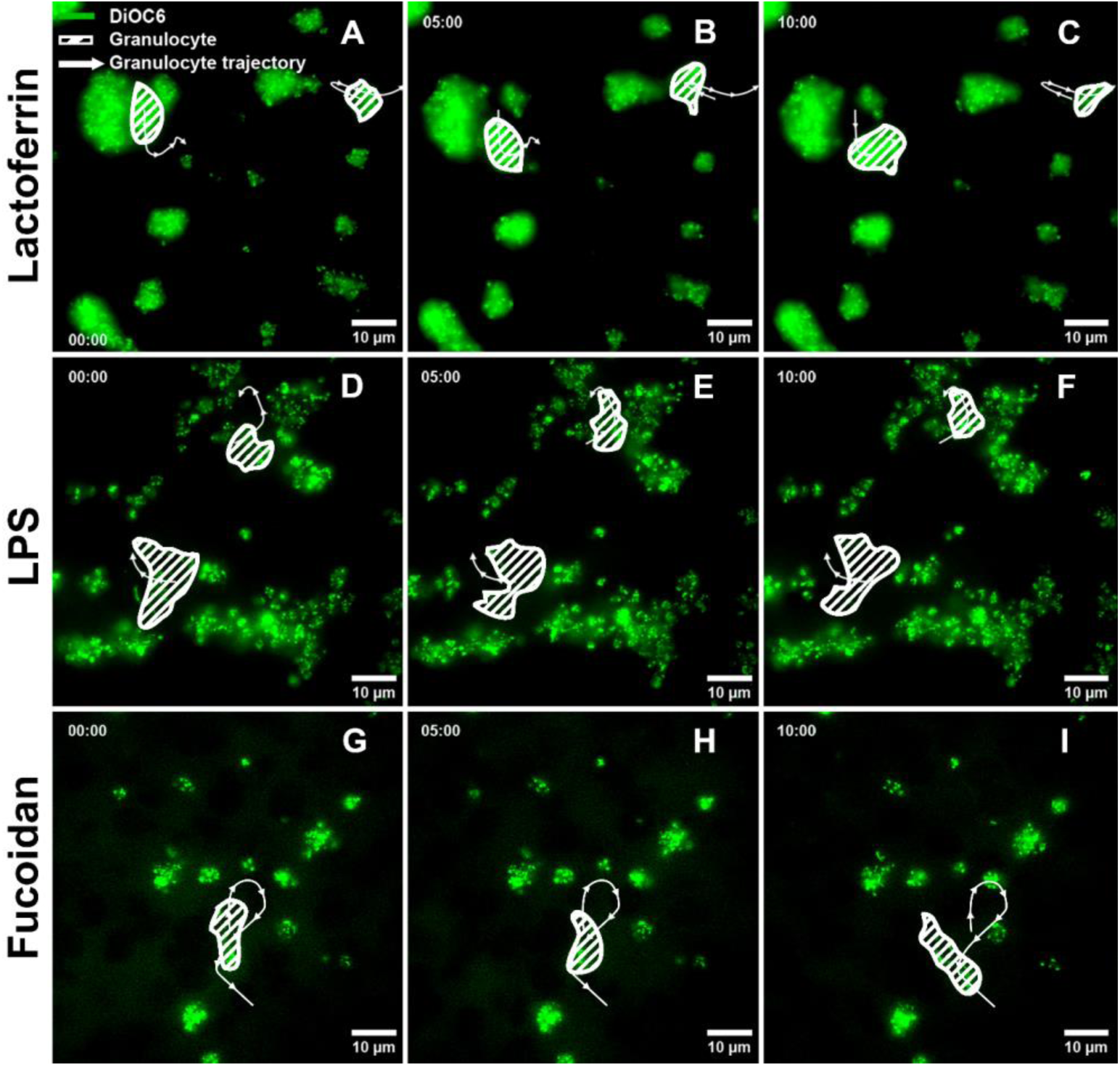
Granulocyte movement around thrombi. Pre-incubated with leukocyte activators anticoagulated whole blood. DIC/epiflourescence (DiOC6 stain) modes were used. Images presented were captured on the 15^th^ minute of perfusion. (A-E) Stained with DiOC6 (green) leukocyte and platelet cell membranes; Leukocyte movement trajectories are shown with white arrows. Indices (a-c) indicate the flow of time: a – 15^th^ min after the start of the experiment, b – +5 min, c – +10 min; (A) hirudinated whole blood as control; (B) pre-incubation with MPO (80 µg/ml); (C) pre-incubation with rLf (200 nM); (D) pre-incubation with LPS (10 µg/ml); (E) pre- incubation with fucoidan (100 µg/ml);

**Figure S4.**
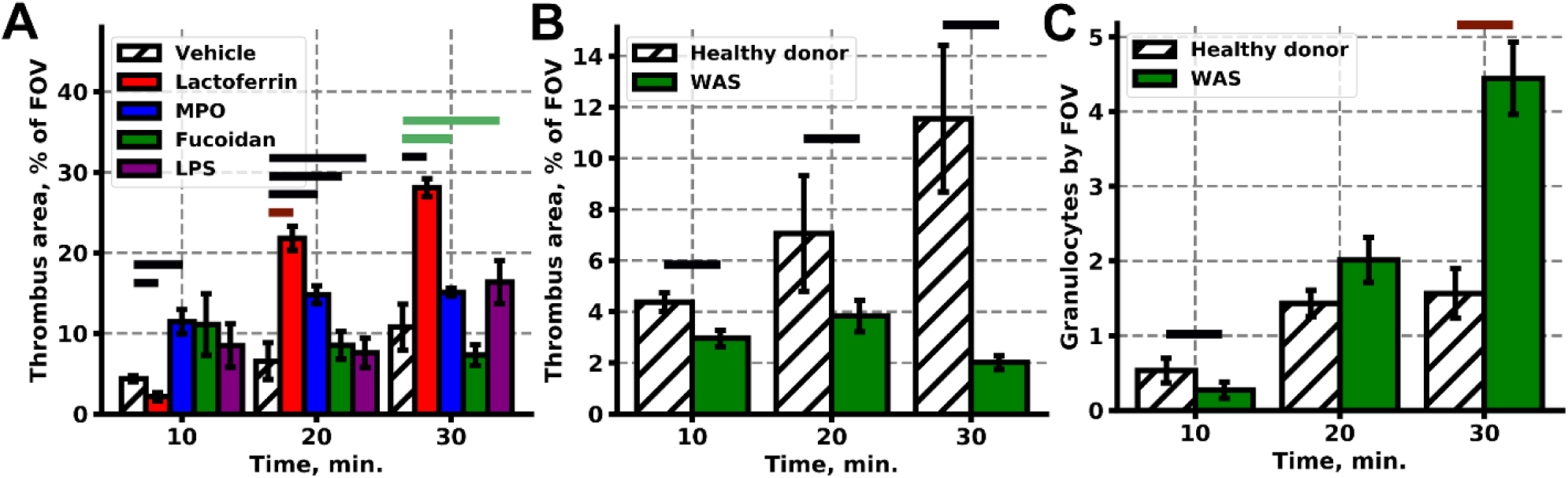
Thrombus area and percentage of highly activated leukocytes in the presence of leukocyte activators and in patients with WAS. (A) - Changes in thrombus area with the flow of time under the influence of blood pre-incubation with leukocytes activators; (B) - Changes in thrombus area with the flow of time in healthy pediatric donors and patients with WAS; (C) - percentage of highly spread leukocytes in patients with WAS is significantly higher from the start of the experiment up to 70% on the 20^th^ minute. Statistical significance was calculated with Mann-Wittney test, green lines indicate p < 0.05, red lines indicate p < 0.01, black lines indicate p < 0.001.

**Figure S5.**
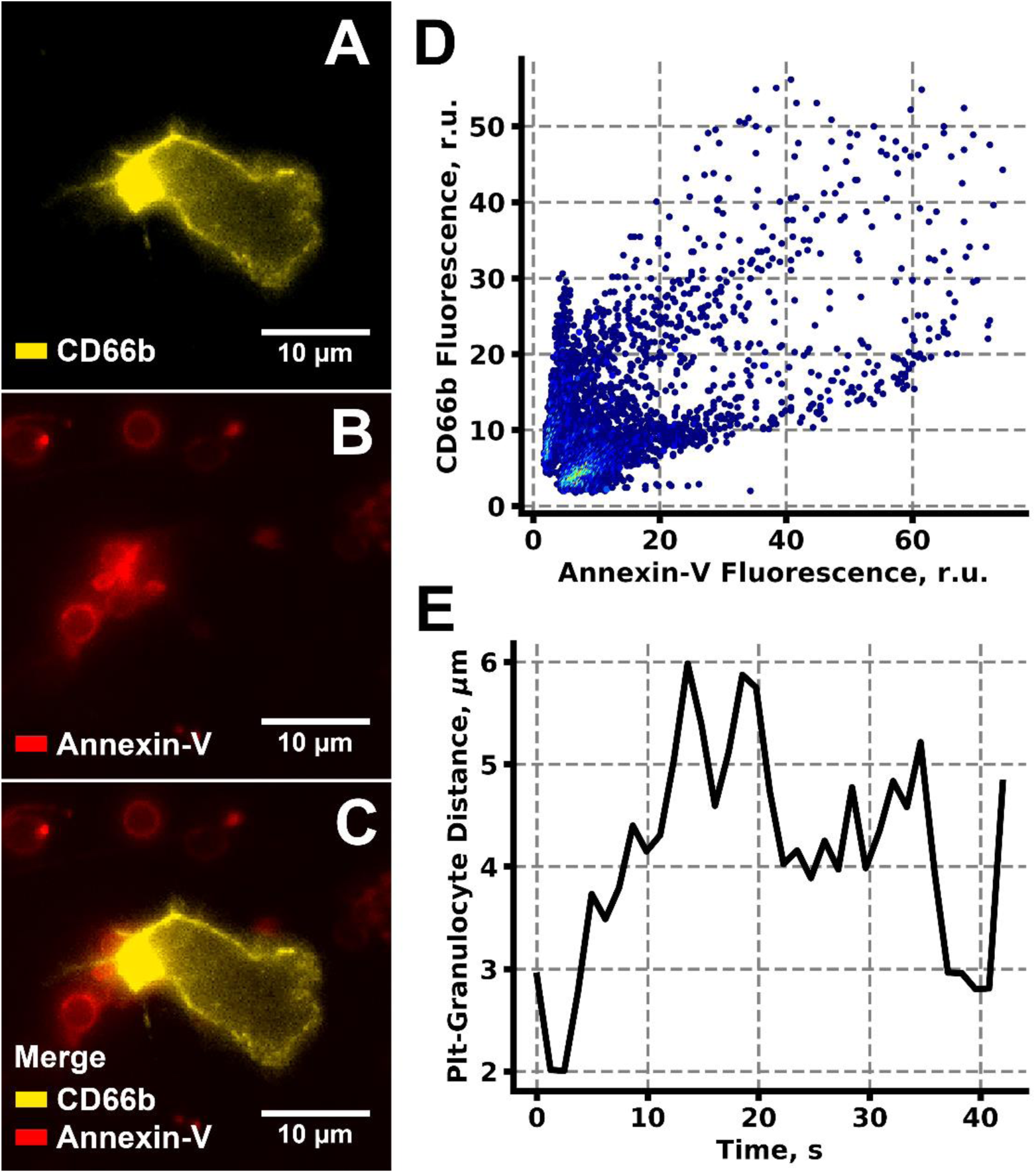
(A-C) CD66b (A) and Annexin (B) staining colocalized (C) in a manner suggesting that granulocytes bear Annexin-V positive platelets. (D) Scatterplot of the Annexin-V and CD66b fluorescence intensity correlation from (C). (E) Distance between the crawling granulocyte and Annexin-V positive platelet from Fig. 4J-L.

## Code listing

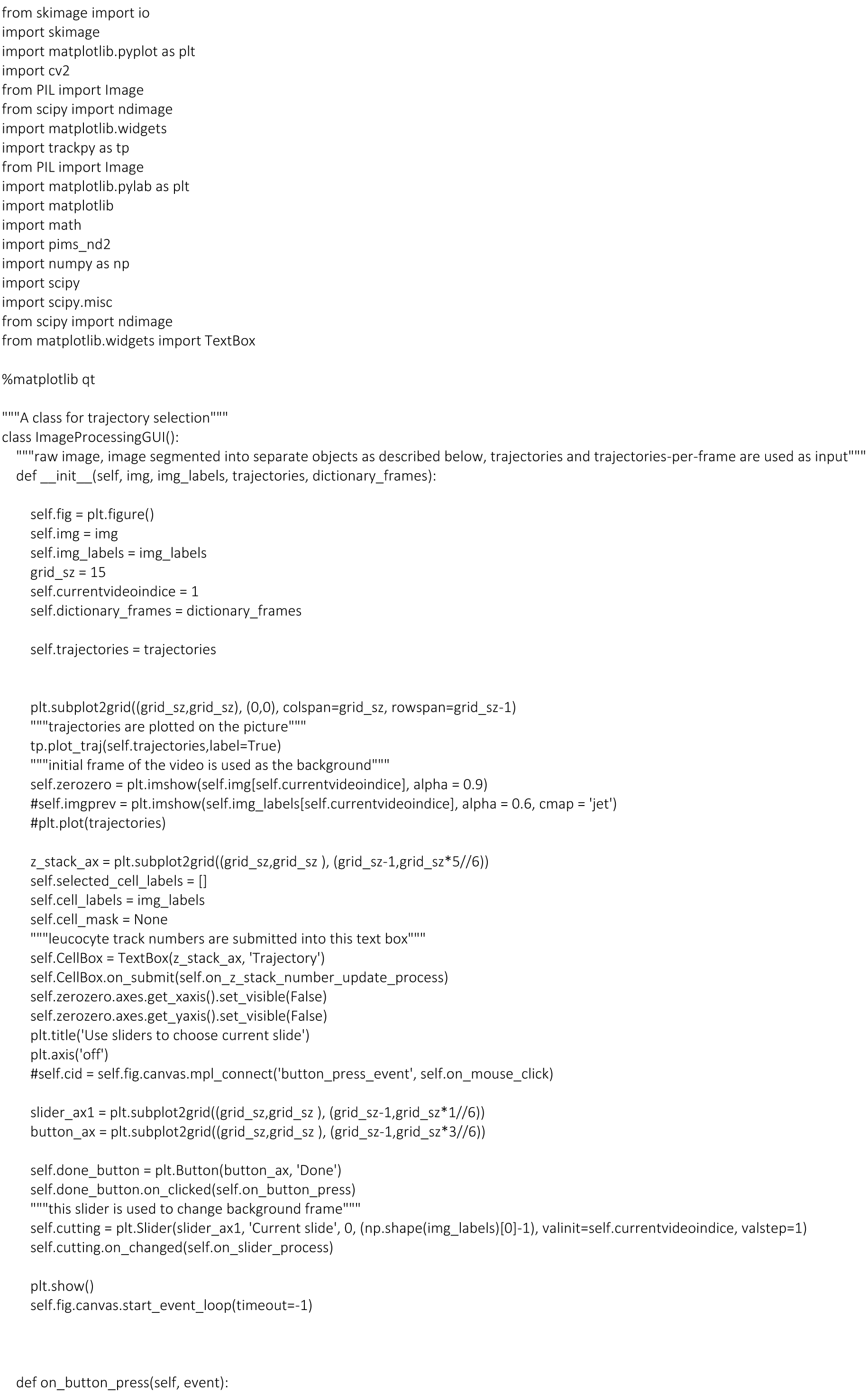
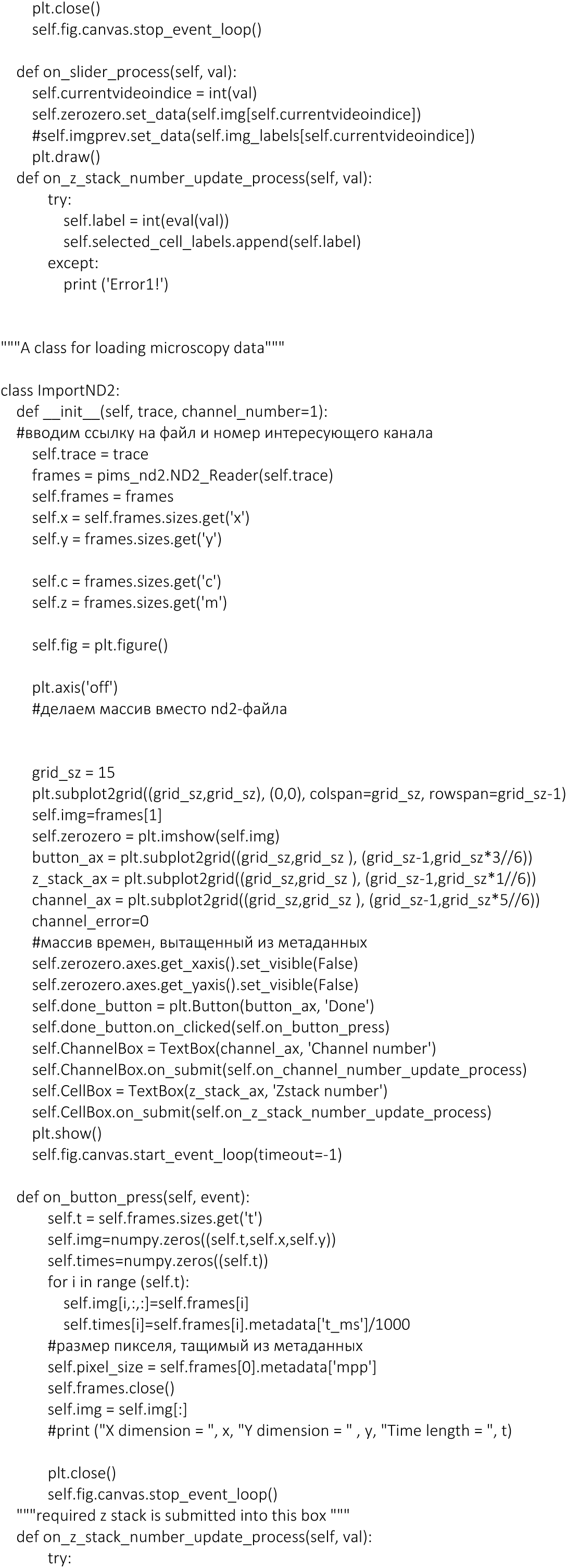
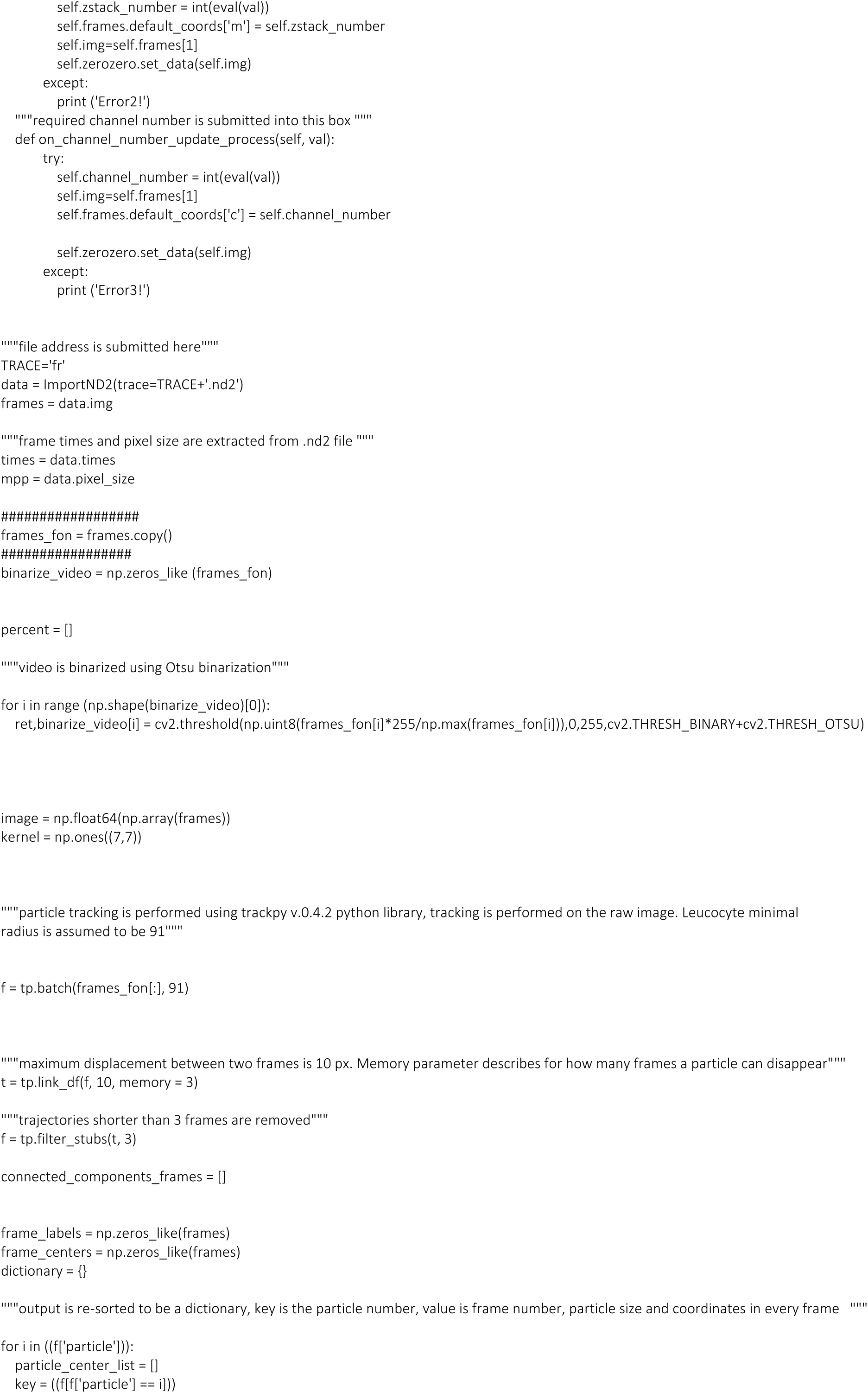
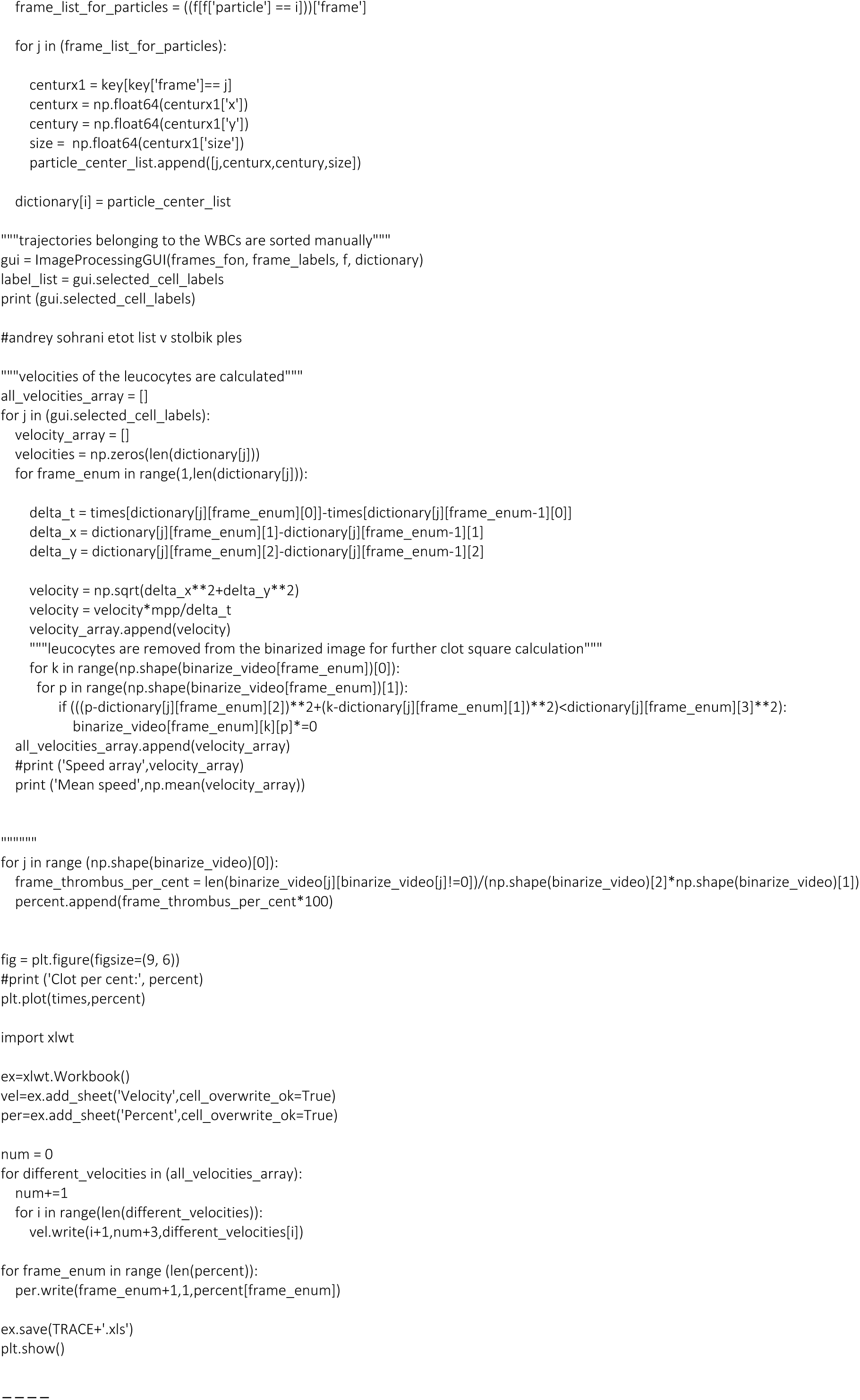

## Notes

### Competing Interest Statement

The authors have declared no competing interest.

